# Coordination of asparagine uptake and asparagine synthetase expression is required for T cell activation

**DOI:** 10.1101/2020.02.28.969774

**Authors:** Helen Carrasco Hope, Rebecca J. Brownlie, Lynette Steele, Robert J. Salmond

## Abstract

T cell receptor triggering by antigen results in metabolic reprogramming that, in turn, facilitates T cells’ exit from quiescence. The increased nutrient requirements of activated lymphocytes are met in part by upregulation of cell surface transporters and enhanced uptake of amino acids, fatty acids and glucose from the environment. However, the role of intracellular pathways of amino acid biosynthesis in T cell activation is relatively unexplored. Asparagine (Asn) is a non-essential amino acid that can be synthesized intracellularly through the glutamine-hydrolyzing enzyme asparagine synthetase (ASNS). We set out to define the requirements for uptake of extracellular Asn and ASNS activity in CD8^+^ T cell activation. At early timepoints of activation, T cells expressed little or no ASNS and, as a consequence, viability and TCR-stimulated growth, activation and metabolic reprogramming were substantially impaired under conditions of Asn deprivation. At later timepoints (>48h of activation), TCR-induced mTOR-dependent signals resulted in upregulation of ASNS, that endowed T cells with the capacity to function independently of extracellular Asn. Thus, we have determined that the coordinated upregulation of ASNS expression and uptake of extracellular Asn is required for optimal T cell effector responses.

## Introduction

Naïve T lymphocytes are small quiescent cells that take up low levels of nutrients to sustain a basal catabolic metabolism. By contrast, antigenic stimulation of T cells results in the upregulation of nutrient receptors and the engagement of anabolic processes in order to generate the biomolecules required for rapid growth, proliferation and effector function. In recent years, critical roles in T cell activation and differentiation have been determined for upregulated expression of the heterodimeric neutral amino acid transporter Slc7a5/Slc3a2 and glutamine transporter Slc1a5 (1, 2). Furthermore, uptake of a number of amino acids, including glutamine (2, 3), leucine (4, 5), tryptophan (6) and methionine (7), has been determined to play important functions in T cell activation. Despite the fact that a number of key amino acids such as glutamine, serine and alanine can be synthesized intracellularly, extracellular sources of these amino acids are required for T cells to meet their metabolic requirements upon exit from quiescence and to facilitate activation (8, 9).

The uptake and metabolism of amino acids plays a key role in numerous intracellular processes in lymphocytes. For example, glutamine (Gln) is used as a building block for protein synthesis, to maintain activation of the mechanistic target of rapamycin complex 1 (mTORC1) (2), to replenish Krebs’ cycle intermediates via glutaminolysis (10), to synthesize glutathione (11) and to fuel post-translational O-GlcNAcylation of key proteins (12). A further potential fate for Gln is the production of asparagine (Asn), via the activity of the Gln-hydrolyzing enzyme asparagine synthetase (ASNS) (13). Interestingly, Asn metabolism and uptake has emerged as a key target in cancer therapy. Thus, ASNS expression is a predictive biomarker in several human cancer types (14, 15) whilst asparaginases, that degrade extracellular Asn, are a key chemotherapeutic drug for the treatment of leukaemia and lymphomas that lack ASNS expression (13). Furthermore, upregulation of ASNS expression via the stress-induced activating transcription factor 4 (ATF4) occurs in B and T cells in response to glucose and amino acid deprivation (16, 17). However, to date, the physiological role of intracellular Asn production via ASNS in T cell responses is unknown. Bacterial asparaginases act to suppress T cell activation (18, 19), implying that T cells require extracellular sources of Asn. However, asparaginases frequently have glutaminase activity and asparaginases that lack this function are less immunosuppressive in mice (20).

In the current work, we set out to determine roles for extracellular Asn uptake and intracellular Asn production via ASNS in T cell activation. We show that in naïve mouse CD8^+^ T cells, ASNS expression is very low or absent, and Asn uptake from the extracellular environment is required for maintenance of cell viability, initial stages of T cell growth and activation. Furthermore, under conditions of Asn deprivation, T cell metabolic switching is compromised resulting in decrease nutrient uptake, reduced glycolytic flux, oxidative phosphorylation and ATP production. T cell receptor triggering and metabolic reprogramming via mTORC1 activation is accompanied by increased expression of ASNS and subsequently T cells can function independently of extracellular sources of Asn. Using gene-trap mice expressing a hypomorphic *Asns* allele, we determined that ASNS expression is essential for effector T cell responses in the absence of extracellular Asn. Together, these data define an essential role for coordinated Asn uptake and biosynthesis in T cell activation and effector responses.

## Results

### Asn uptake is required for initial cell growth, survival and activation following TCR stimulation

We tested the impact of Asn and/or Gln deprivation on TCR-induced cell growth and activation. Naïve TCR transgenic OT-1 CD8^+^ T cells were stimulated for 24h with cognate peptide SIINFEKL in Dulbecco’s Modified Eagle’s Medium (DMEM) supplemented ± Asn ± Gln. Deprivation of either Asn or Gln alone reduced, whilst the combined absence of both amino acids substantially impaired T cell viability (Figure 1A). The majority (70-80%) of T cells cultured in the presence of both amino acids had initiated blastogenesis following 24h of TCR stimulation (Figure 1B). By contrast, OT-1 T cells stimulated with peptide in the absence of Gln or the combined absence of Gln and Asn did not form a blast cell population (Figure 1B). An intermediate phenotype was determined for OT-1 cells activated in the absence of Asn alone; a reduced but reproducible proportion (20-40%) of the cells had a blast phenotype (Figure 1B). A fundamental role for amino acids is to act as building blocks for protein synthesis. Consistent with the impact on T cell growth, we determined that levels of protein synthesis, as measured by assessment of uptake and incorporation of O-propargyl-puromycin (OPP), by TCR-stimulated OT-1 T cells were substantially reduced in the absence of extracellular Asn and/or Gln (Figure 1, C and D).

**Figure 1:**
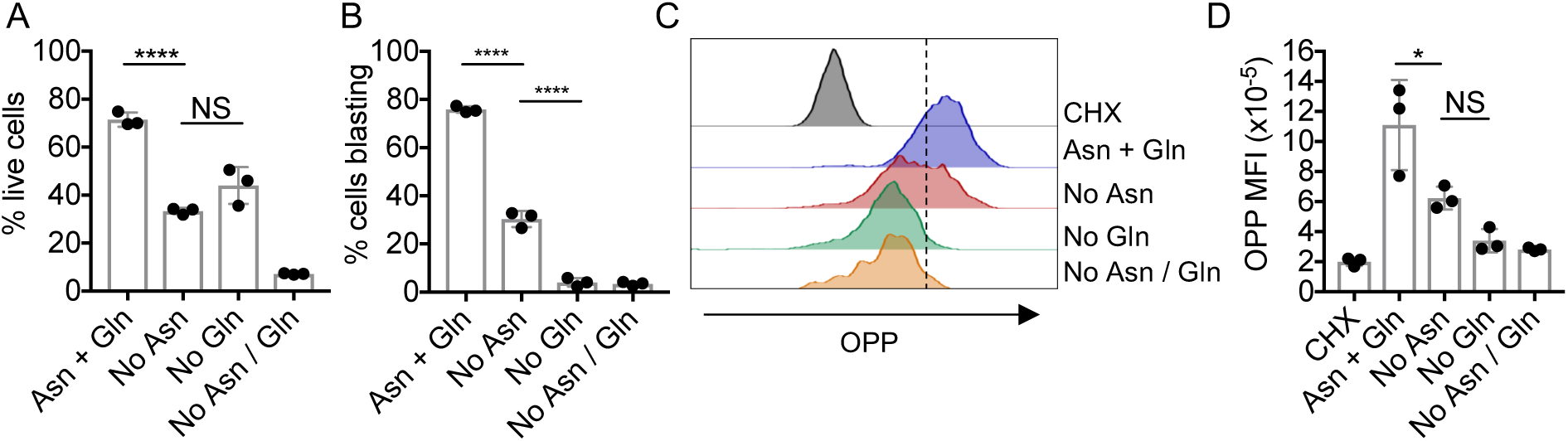
Extracellular Asn is required to maintain viability and initiate cell growth following TCR stimulation. OT-1 TCR transgenic T cells were stimulated with cognate SIINFEKL peptide for 24h in DMEM supplemented ± Asn ± Gln, as indicated. (A) T cell viability was assessed by exclusion of Live/Dead Aqua dyes and FACS. (B) T cell growth was assessed by analysis of forward scatter and side scatter area (FSC-A/SSC-A) parameters on gated live cells by flow cytometry. (C) Nascent protein synthesis was assessed by incorporation of OPP, intracellular staining and labelling using Click chemistry reagents and FACS analysis. Cycloheximide (CHX) was used as a negative control. (D) OPP mean fluorescence intensity (MFI) was assessed by FACS analysis. Data are from one of two or more repeated experiments. NS – not significant, * - p<0.05, **** - p<0.0001 as assessed by one-way ANOVA, with Tukey’s multiple comparisons test.

Next, we assessed the impact of amino acid deprivation on TCR-induced activation marker and cytokine expression. Antigen-stimulated OT-1 T cell IL-2 secretion was inhibited 70-90% by deprivation of Asn and/or Gln (Figure 2A). Furthermore, whilst TCR-induced upregulation of the very early activation marker CD69 was unimpeded, cell surface expression of CD25 and transferrin receptor CD71 was partially or completely inhibited by a lack of extracellular Asn and/or Gln (Figure 2B). Similarly, the proportions of T cells expressing key transcription factors eomesodermin (Figure 2C) and Tbet (Figure 2D) were reduced under conditions of Asn or Gln deprivation. TCR-induced expression of cytolytic effector protein granzyme B was low under all media conditions after 24h of stimulation. Asn-deprivation impaired TCR-induced upregulation of granzyme B, as assessed following 48h of stimulation (Figure 2E). Both proportions of positive cells (Figure 2F) and levels of granzyme B within positive cells, as assessed by mean fluorescence intensity (Figure 2G), were reduced compared to control conditions. As compared to either Asn deprivation or control conditions, Gln deprivation more severely impaired TCR-induced granzyme B expression (Figure 2, E-G).

**Figure 2:**
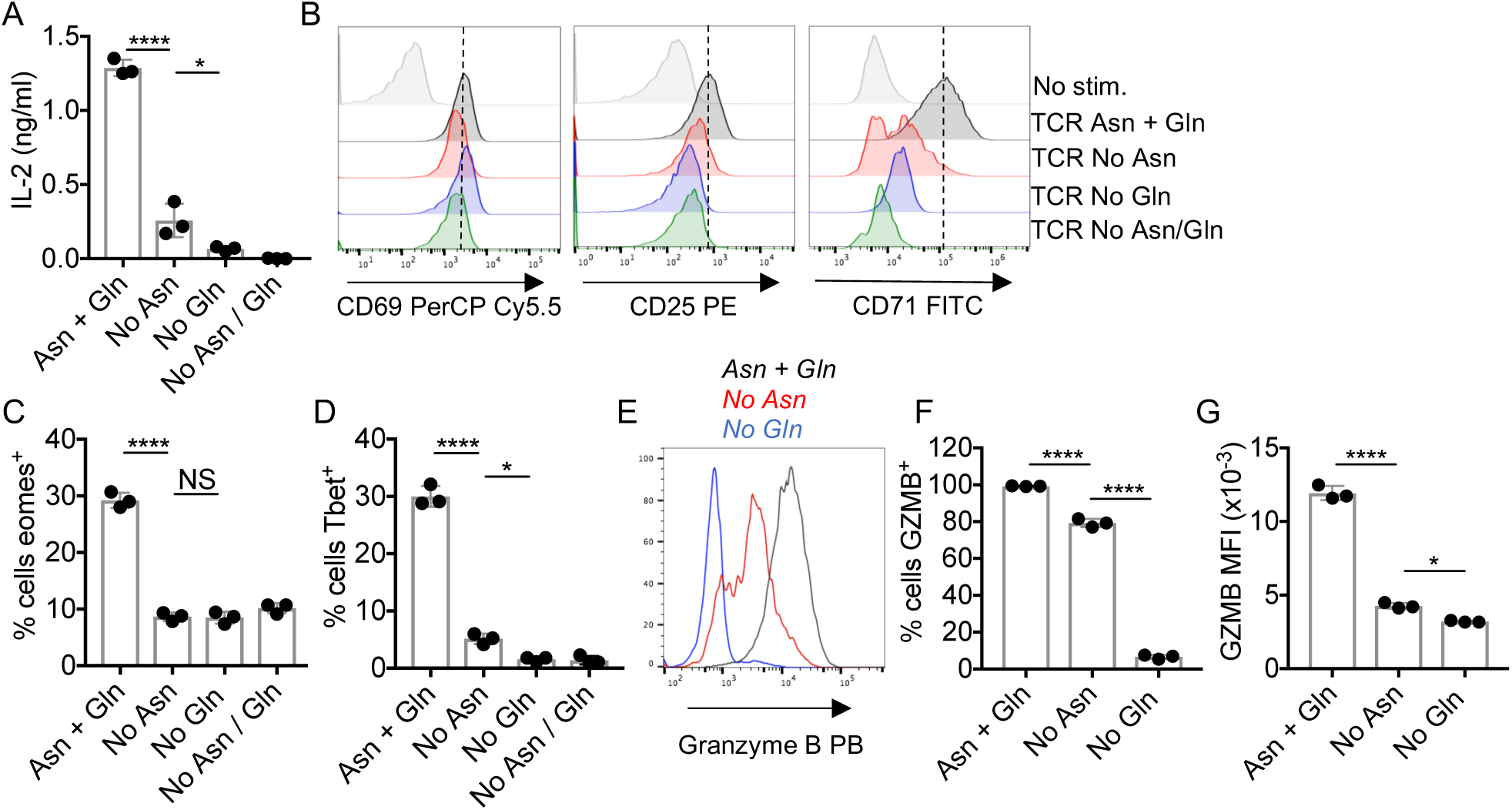
Extracellular Asn is limiting for initial TCR-induced T cell activation. OT-1 TCR transgenic T cells were stimulated with cognate SIINFEKL peptide for 24h (A-D) or 48h (E-G) in DMEM supplemented ± Asn and/or Gln, as indicated. (A) Levels of IL-2 in culture supernatants were assessed by ELISA. (B) Histograms show levels of cell surface activation marker expression on live-gated cells. Data are representative of one of at least three repeated experiments. Proportions of live-gated cells expressing transcriptions factors eomesodermin (eomes) (C) and Tbet (D) were determined by intracellular staining and FACS. (E) Representative histograms show levels of intracellular granzyme B in live-gated cells. Proportions of granzyme B (GZMB)-positive cells (F) were determined. Relative levels of expression of granzyme B in live granzyme B^+^ cells were assessed as mean fluorescence intensity (MFI) (G). All data are from one of at least 3 repeated experiments. NS – not significant, * - p<0.05, **** - p<0.0001 as assessed by one-way ANOVA, with Tukey’s multiple comparisons test.

IL-2 is a potent T cell growth factor and mitogen. Thus, it was possible that a failure to produce IL-2 was responsible for reduced OT-1 T cell viability, growth and activation under conditions of amino acid deprivation. However, addition of exogenous recombinant IL-2 did not rescue T cell viability, protein synthesis, or enhance TCR-induced expression of granzyme B in Asn- or Gln-deprived media (Supplemental Figure 1). Taken together these data indicate that CD8^+^ T cells require extracellular sources of Asn and Gln to maintain cell viability and initiate cell growth, protein synthesis and activation following TCR stimulation. Either amino acid alone is sufficient to maintain a reduced proportion of viable cells, however the combined absence of Asn and Gln results in catastrophic levels of cell death.

### Asn availability regulates TCR-induced metabolic reprogramming

Nutrient uptake and metabolic reprogramming following TCR-stimulation is highly coordinated. Therefore, it was of interest to determine whether the availability of Asn impinged upon T cell metabolism. The tryptophan metabolite kynurenine is transported into T cells via the neutral amino acid transporter Slc7a5/CD98 (21), that also serves as a transporter for essential amino acids tryptophan, leucine, phenylalanine and methionine. We took advantage of the fluorescent properties of kynurenine as well as the glucose analog 2-NB-d-glucose and long chain fatty acid analog BODIPY™ C16 to assess the impact of amino acid deprivation on TCR-induced nutrient uptake. FACS analysis demonstrated that OT-1 T cells activated for 24h in the absence of Asn and/or Gln had reduced capacity to uptake glucose (Figure 3A), kynurenine (Figure 3B) and BODIPY™ C16 (Figure 3C) as compared to cells activated in the presence of both amino acids. Reduced TCR-induced nutrient uptake was associated with impaired cell surface expression of nutrient transporters GLUT1/Slc2a1, CD98 and CD36 under conditions of Asn ± Gln deprivation (Figure 3, D-F).

**Figure 3:**
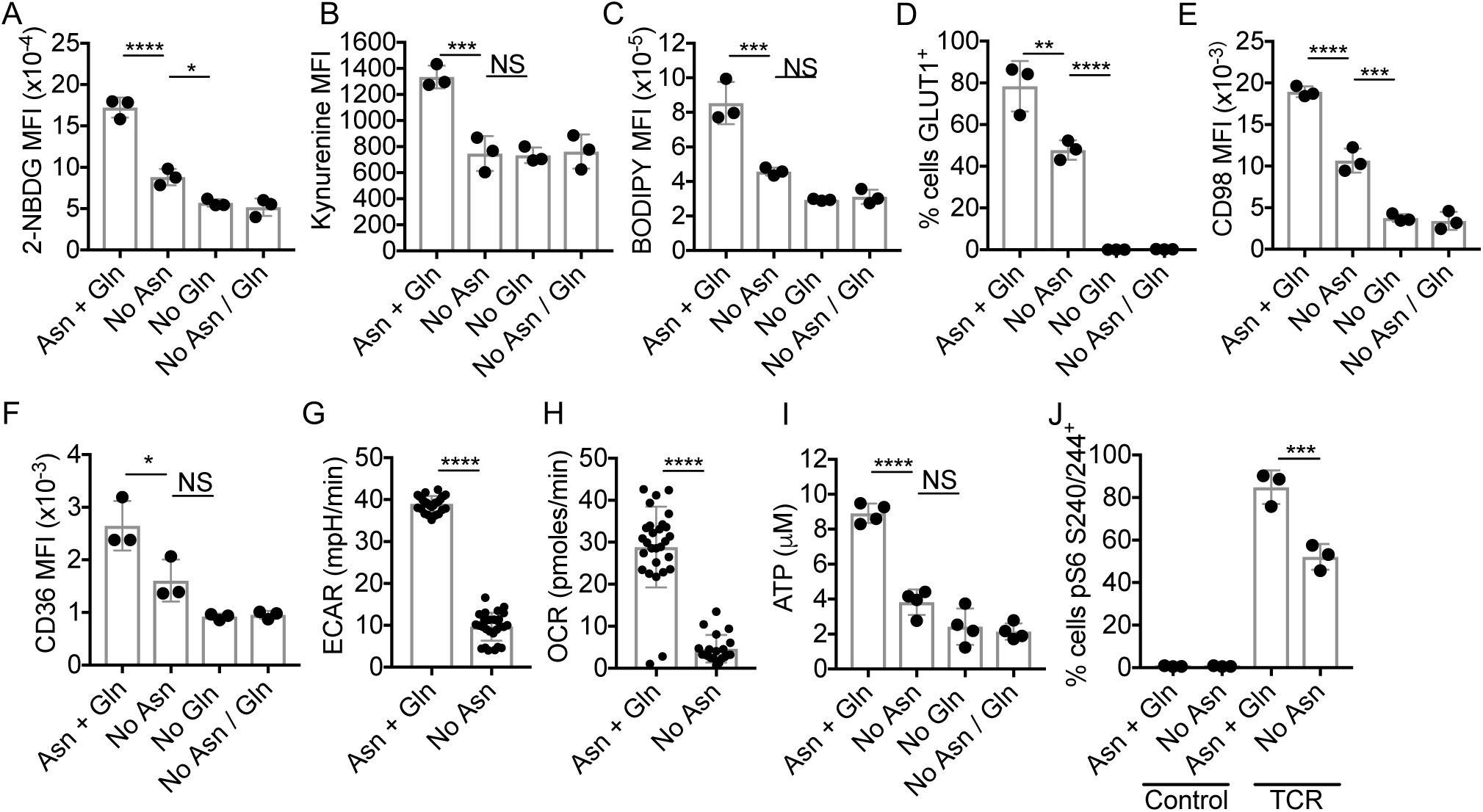
Extracellular Asn is limiting for initial TCR-induced metabolic reprogramming. OT-1 T cells were stimulated with cognate SIINFEKL peptide for 24h in DMEM supplemented ± Asn and/or Gln, as indicated. Uptake of fluorescent glucose analogue 2-NBDG (A), kynurenine (B), and long chain fatty acid BODIPY™-C16 (C), and cell surface expression of transporters GLUT1 (D), CD98 (E) and CD36 (F) were assessed by FACS analysis. Data are mean fluorescence intensity values (MFI) or % cells positive, as indicated. (G, H). Extracellular acidification rate (ECAR) and oxygen consumption rate (OCR) of activated OT-1 T cells were assessed using a Seahorse metabolic analyser. (I) ATP production was assessed using a luminescence-based assay and luminometer. (J) Proportions of phospho-ribosomal protein S6 (pS6 S240/244)-positive cells were assessed by intracellular staining and FACS analysis. Data are representative of two (G, H) or three repeated experiments. NS – not significant, * - p<0.05, ** - p<0.01, *** - p<0.001, **** - p<0.0001, as determined by one-way ANOVA with Tukey’s multiple comparisons test (A-F, I), two-tailed Student’s *t*-test (G, H) or two-way ANOVA (J).

Analysis of extracellular acidification rate (ECAR) and oxygen consumption rate (OCR) serve as proxy measures for lactate secretion / glycolytic rate and oxidative phosphorylation, respectively. Seahorse metabolic analysis demonstrated that OT-1 T cells activated under conditions of Asn deprivation had reduced ECAR (Figure 3G) and OCR (Figure 3H), suggestive of overall reduced levels of cellular metabolism. Consistent with these data, levels of OT-1 T cell ATP production were reduced by 50-75% in Asn and/or Gln-deprived media (Figure 3I).

The availability of amino acids, including Gln, has previously been shown to play a central role in T cell activation and metabolism through regulation of the activity of the mTORC1 kinase complex (1, 2). Analysis of phosphorylation of the downstream substrate ribosomal protein S6 (22) determined that, in the absence of extracellular Asn, TCR-induced mTORC1 activation was impaired but not blocked (Figure 3J). Together these data indicate that, at initial stages of T cell activation, the bioavailability of Asn determines the extent of TCR-induced nutrient uptake, glycolytic flux, OXPHOS and energy production, with a partial effect on mTORC1 activation.

### T cells lose requirement for extracellular Asn upon prolonged TCR stimulation

It was noteworthy that at early timepoints (24h), in contrast to Gln deprivation, a reduced but consistent proportion of T cells stimulated in the absence of extracellular Asn demonstrated activation profiles comparable to cells stimulated under control conditions (i.e. blast phenotype, elevated activation marker expression/OPP incorporation, phospho-S6^+^) (Figures 1-3). We sought to determine whether T cell activation remained dependent upon extracellular Asn at later timepoints. Following 72h of peptide stimulation, levels of OT-1 cell viability were comparable in Asn-deprived as compared to control media conditions (Figure 4A). By contrast, OT-1 T cell viability was much impaired under conditions of Gln-deprivation (10-20%), whilst T cells did not survive prolonged culture in the combined absence of both amino acids (Figure 4A). Viable T cells activated for 72h under control conditions and in the absence of Asn, but not Gln, were uniformly large blasting cells as determined by analysis of FSC/SSC parameters (Figure 4B). Furthermore, analysis of Cell Trace Violet dilution after 72h of activation indicated that Asn-deprivation reduced but did not block antigen-stimulated T cell proliferation (Figure 4, C and D). By contrast, Gln deprivation completely prevented T cell proliferation *in vitro*.

**Figure 4:**
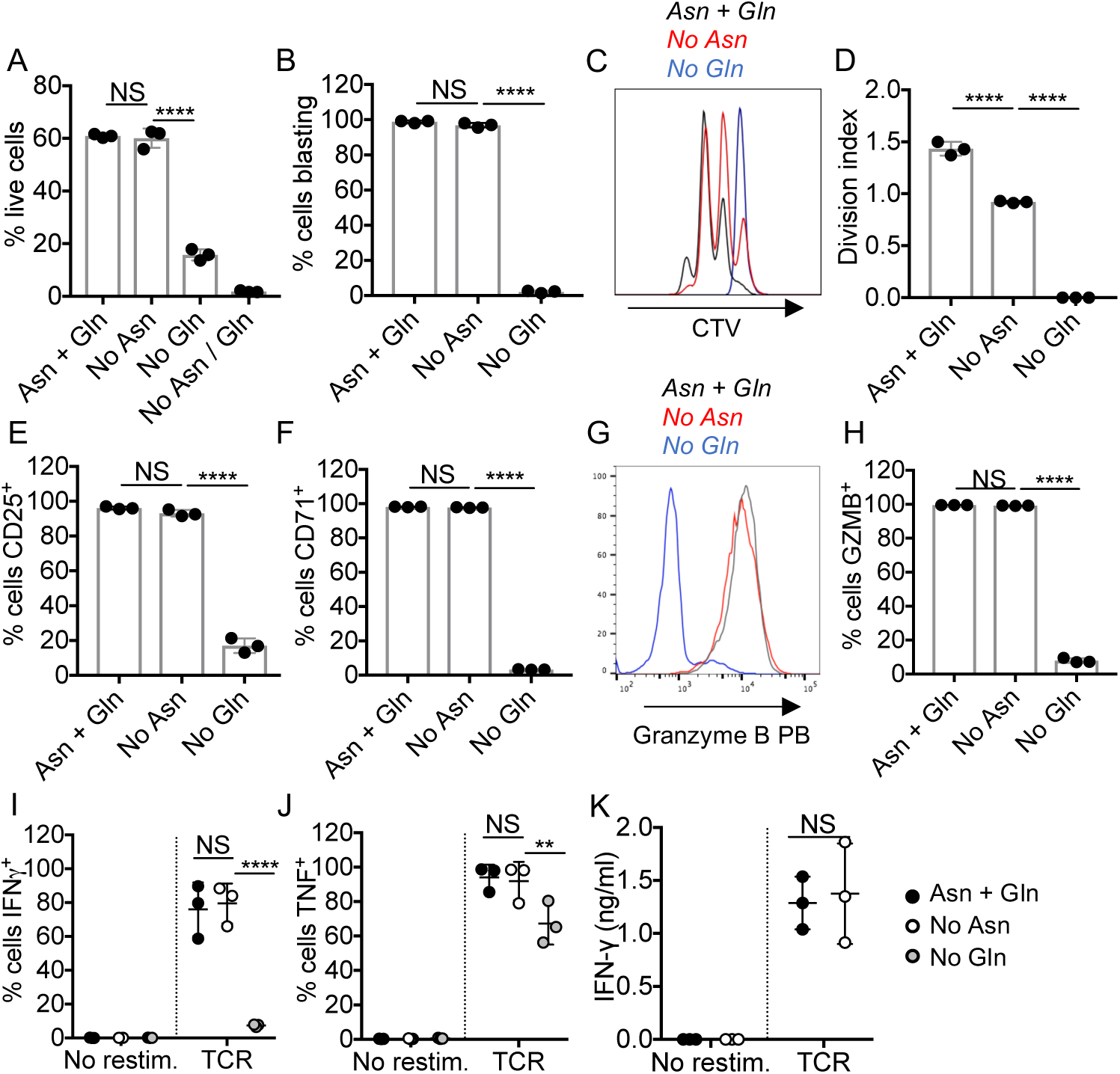
T cells lose requirement for extracellular Asn upon prolonged activation. OT-1 T cells were stimulated with cognate SIINFEKL peptide for 72h (A-G) or were differentiated for 6d (H, I) in DMEM supplemented ± Asn and/or Gln or complete IMDM (J, K), as indicated. T cell viability (A) and proportions of blasting cell populations (B) were assessed by exclusion of Live/Dead Aqua dyes and analysis of forward scatter and side scatter area (FSC-A/SSC-A) parameters by flow cytometry, respectively. For analysis of TCR-induced proliferation, cells were labelled with Cell Trace Violet prior to stimulation. Histograms are representative of data from 3 repeated experiments (C). Division indices were calculated using FlowJo software (D). Proportions of live-gated T cells expressing CD25 (E), CD71 (F) or granzyme B (G, H) were determined by flow cytometry. (I, J) Effector CTLs were differentiated for 6d in DMEM ± Asn / Gln, then restimulated in complete IMDM for 4h. Proportion of IFNγ- and TNF-positive cells were determined by intracellular staining and FACS analysis. (K) Effector CTLS were differentiated in complete IMDM for 5d, before a 24h washout in DMEM ± Asn, then were restimulated in DMEM ± Asn for 24h. Levels of IFNγ in supernatants were determined by ELISA. Data are representative of 2 (C, D) or at least 3 repeated experiments. NS – not significant, ** - p<0.01, **** - p<0.0001, as determined by one-way ANOVA with Tukey’s multiple comparisons test (A, B, D-F, H) or or two-way ANOVA with Sidak’s multiple comparisons test (I-K).

Comparison of OT-1 T cell phenotypes after 72h of peptide stimulation indicated that levels of CD25, CD71 and granzyme B expression were equivalent in control cells and cells activated in Asn-deprived conditions (Figure 4, E-H). By contrast, T cells stimulated in the absence of Gln had very low levels or absent expression of cell surface activation markers and granzyme B (Figure 4, E-H). Furthermore, and in contrast to results from earlier timepoints, OT-1 T cells activated for 72h in the absence of Asn had unimpaired capacity to uptake glucose and long-chain fatty acids (Supplemental Figure 2, A and B), whilst levels of ATP production were comparable to control levels (Supplemental Figure 2C).

We assessed the functional capacity of effector CTLs differentiated for 6 days in the absence of either Asn or Gln. OT-1 T cells differentiated in the absence of Gln did not produce IFNγ and reduced proportions of cells were competent to produce TNF upon antigenic re-stimulation in nutrient-replete media (Figure 4, I and J). By contrast, CTLs differentiated in the absence of Asn produced equivalent levels of effector cytokines upon re-stimulation in nutrient-replete media as compared to control cells (Figure 4, I and J). These data indicate that extracellular Gln but not Asn is essential for CD8^+^ T cell differentiation to an inflammatory phenotype. We wished to test whether there was any requirement for extracellular sources of Asn in effector T cell function. To this end, we differentiated OT-1 T cells in nutrient-replete IMDM media for 5 days, transferred to DMEM supplemented with Asn ± Gln overnight prior to re-stimulation of the resultant CTLs. Asn-deprivation did not impede effector T cell secretion of IFNγ (Figure 4K).

### T cells upregulate expression of ASNS via the mTOR signalling pathway

We reasoned that the initial dependence of T cells on extracellular Asn could be a consequence of low/absent ASNS expression, and that ASNS might be upregulated following T cell activation. To test this, OT-1 T cells were activated for various time-periods in nutrient replete conditions and western blot analysis of ASNS expression performed. The data indicated that ASNS was not expressed in naïve OT-1 cells and at very low levels in thymocytes, but was upregulated in response to TCR stimulation (Figure 5A). Activation of the mTOR pathway is central to metabolic reprogramming following T cell activation (23), whilst our data suggested that Asn availability was rate-limiting for mTOR activation at early stages of T cell activation (Figure 3). Western blots showed that treatment with rapamycin blocked TCR-induced upregulation of ASNS (Figure 5A), suggesting that ASNS upregulation is a feature of mTORC1-dependent T cell metabolic reprogramming.

**Figure 5:**
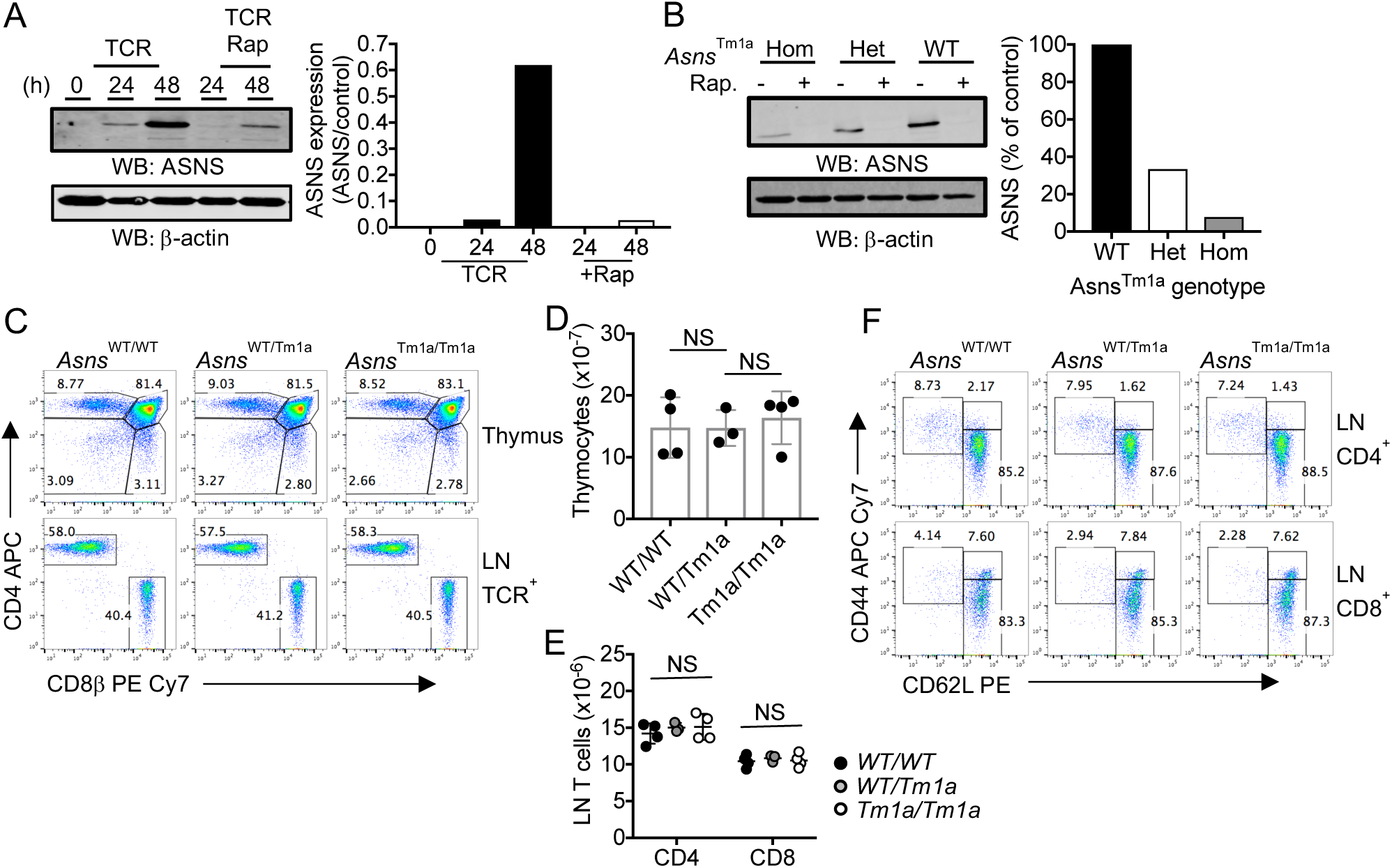
T cell development and homeostasis is unimpaired in ASNS-deficient mice. (A) Western blots show levels of ASNS protein in lysates of OT-1 T cells stimulated in IMDM with SIINFEKL peptide ± rapamycin (rap), for the indicated time periods, or thymocytes. (B) ASNS levels in lysates from activated lymph node T cell from homozygous, heterozygous or wild-type (WT) *Asns*^Tm1a^ mice. For (A, B) β actin serves as a protein loading control. Values in bar graph represent relative ASNS expression levels (ASNS/Actin) calculated using the LI-COR Odyssey imaging system from 1 of 2 repeated experiments. (C) Thymocyte and lymph node (LN) cell populations from littermate Asns^Tm1a^ mice were analysed by flow cytometry. Total thymocytes (D) and LN CD4 and CD8 T cells (E) were enumerated from 3-4 age-matched male mice of each genotype. (F) Proportions of naïve, effector and memory cells within gated CD4^+^ and CD8^+^T cell populations were analysed by analysis of CD44 and CD62L expression by flow cytometry. FACS dotplots are representative of 5-6 mice of each genotype. Values on dotplots represent % cells within the defined gates. NS – not significant, as determined by one-way ANOVA.

### T cell development and homeostasis is unimpaired in ASNS gene-trap mice

In order to determine the role of ASNS expression in T cells, we assessed *Asns*^Tm1a^ mice, a genetically-modified mouse strain with a gene-trap insertion in intron 2 of the *Asns* locus. Previous studies of the role of ASNS in neural development determined that the gene-trap approach resulted in a hypomorphic *Asns* allele (24). Western blotting analysis of activated lymph node T cell lysates determined that homozygous expression of the hypomorphic allele resulted in a >90% decrease in ASNS protein expression, as compared to wild-type littermate controls, whilst heterozygous cells expressed intermediate levels (Figure 5B). For all genotypes, TCR-induced ASNS expression was abolished by rapamycin (Figure 5B), consistent with the results from OT-1 TCR transgenic experiments (Figure 5A). We wished to determine whether ASNS-deficiency impacted upon T cell developmental processes. Flow cytometry analysis of thymus and lymph node cells determined that homozygous *Asns*^Tm1a^ mice had similar proportions and numbers of thymocyte populations and mature CD4^+^ and CD8^+^ T cells as compared with wild-type and heterozygous controls (Figure 5, C-E). Furthermore, proportions of naïve (CD44^low^ CD62L^high^), central memory (CD44^high^CD62L^high^) and effector/effector memory (CD44^high^CD62L^low^) phenotype CD4^+^ and CD8^+^ T cells were similar in *Asns*^Tm1a^ and control mice (Figure 5F). These data indicate that homozygous expression of the hypomorphic *Asns* allele does not impinge on T cell development in the thymus nor on peripheral T cell homeostasis, in a specific-pathogen free animal facility.

### ASNS expression is required for T cell activation in the absence of extracellular Asn

To test the role for ASNS in T cell activation, we stimulated polyclonal lymph node cells from *Asns*^*Tm1a*^ and control mice for 48h with plate-bound CD3/CD28 antibodies in DMEM media supplemented ± Gln and/or Asn. Similar to results using OT-1 T cells, FACS analysis of polyclonal CD8^+^ T cells demonstrated impaired T cell viability in Asn- and Gln-deprived conditions (Figure 6A). Interestingly, ASNS-deficient T cells had lower cell viability under conditions of Asn-deprivation, than control T cells (Figure 6A). Similarly, TCR-induced CD8^+^ T cell blastogenesis was completely abrogated by the combination of genetic ASNS-deficiency and extracellular Asn deprivation (Figure 6, B and C). The combined absence of extracellular Asn and ASNS-deficiency also prevented TCR-induced upregulation of activation marker CD71 and Granzyme B (Figure 6, D and E). Furthermore, population expansion and survival of *Asns*^Tm1a^ CD8^+^ T cells over 6 d of culture in Asn-free media was markedly impaired as compared to control cells (Figure 6F). Taken together, these data indicate that T cell activation, metabolic reprogramming and mTOR-dependent upregulation of ASNS enables T cells to function independently of extracellular sources of Asn.

**Figure 6.**
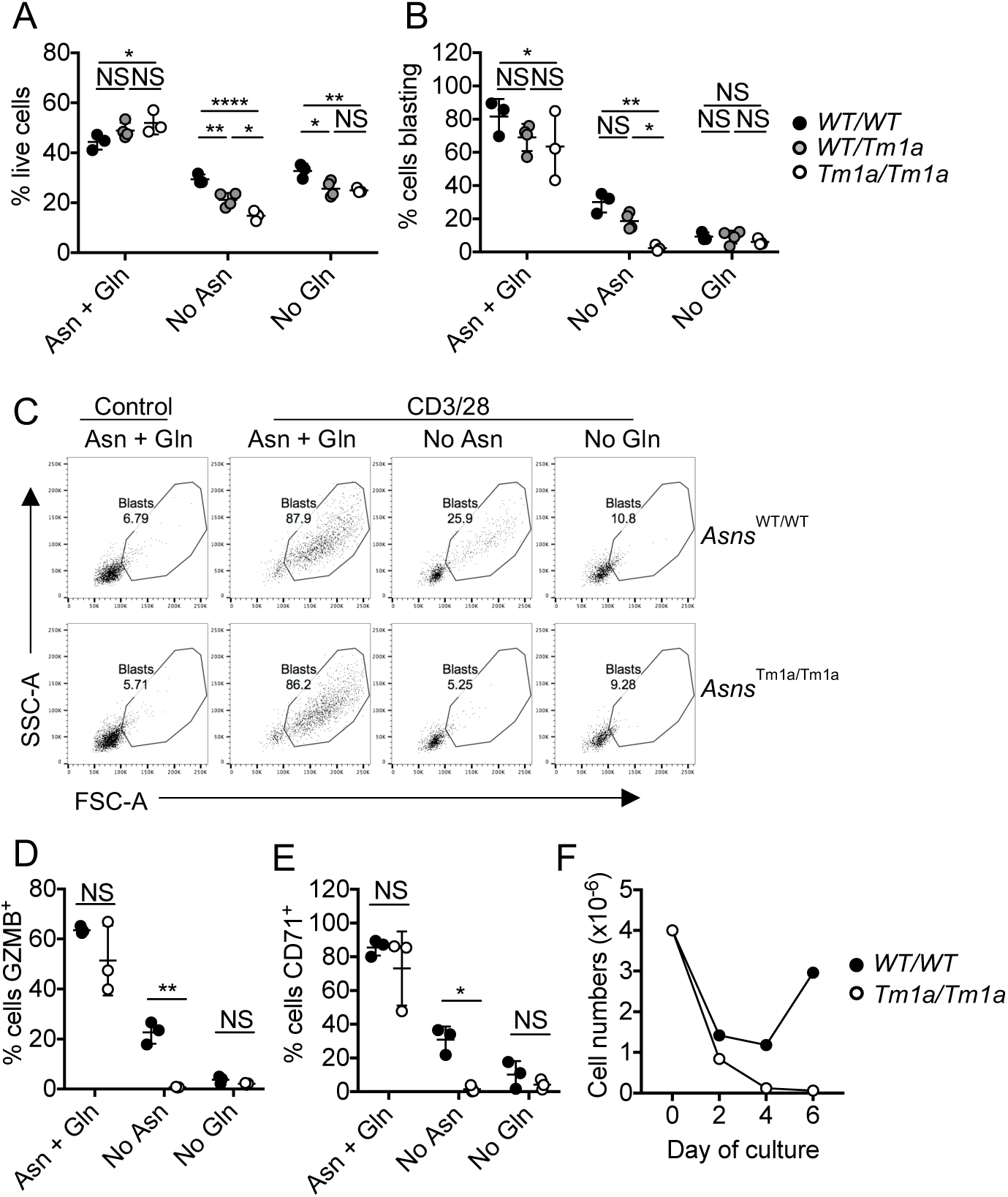
ASNS expression enables T cell responses in Asn-depleted conditions. Lymph node T cells from *Asns*^Tm1a^ mice and controls were stimulated with anti-CD3/28 antibodies for 48h in DMEM ± Asn or Gln. Cell viability was assessed by exclusion of Live-Dead Aqua dye and flow cytometry (A). Proportions of gated live CD8^+^ T cells undergoing blastogenesis were assessed by FACS analysis of FSC-A/SSC-A parameters (B, C). Control cells were cultured in the presence of IL-7 that maintains cell viability without inducing T cell activation (C). Gated live CD8^+^ T cells were analysed by FACS for levels of intracellular granzyme B (D) and cell surface CD71 (E). (F) Purified control and *Asns*^Tm1a^ CD8^+^ T cells were activated in Asn-free DMEM for 6 days. Live cells were enumerated throughout the timecourse of the experiment. In all cases data represent one of at least 3 repeated experiments. NS – not significant, * - p< 0.05, ** - p<0.01, **** - p<0.0001, as determined by two-way ANOVA with Sidak’s multiple comparisons tests.

## Discussion

T cell activation is accompanied by, and dependent upon, metabolic reprogramming and substantial upregulation of nutrient uptake. In the current work, we determined that, initially, antigen-stimulated CD8^+^ T cells require extracellular sources of the non-essential amino acid Asn to maintain viability, to grow and to initiate activation and metabolic remodelling. However, upon prolonged stimulation, T cells diverge from a strict Asn-dependent phenotype as a consequence of upregulation of ASNS expression. By contrast, Gln remains an essential amino acid for T cells throughout all stages of activation and differentiation.

The availability of nutrients is key to immune functionality. Glutamine is the most abundant amino acid in human serum, whilst the results of the current work confirm previous studies’ findings that expression of Gln transporters and Gln uptake is critical for T cell activation (2). It had been assumed that lymphocytes are auxotrophic for Asn through the interpretation of studies of the impact of bacterial asparaginases on transformed and non-transformed lymphocyte populations (13, 18, 19). Our results confirm that extracellular Asn is required for initial stages of T cell activation. In the absence of Asn or Gln alone, TCR stimulation results in substantial levels of T cell death. Nonetheless, our data indicate that Asn can partially compensate for the absence of Gln, and vice versa, as in the combined absence of both amino acids, TCR-stimulated T cells fail to survive. Similar results have been reported for transformed cells; in the absence of Gln, Asn is required to maintain viability and proliferation of cancer cells, whilst high ASNS is associated with poor prognosis in several cancer types (25, 26).

Defective T cell activation, growth and metabolism in Asn-deprived conditions was associated with, and likely to be a consequence of, a reduced capacity for protein synthesis. Asn deprivation results in translational pausing at Asn codons and decreased protein synthesis with a dominant effect on Asn-rich proteins (27). In breast cancer cells, Asn availability influences expression of proteins that regulate epithelial-to-mesenchymal transition, with subsequent effects on metastasis (14). Whether reduced Asn availability impacts on specific subsets of proteins or has a more global effect on the T cell proteome remains to be investigated. In addition to its fundamental role in protein synthesis, Asn has been reported to function as an exchange factor for additional amino acids in tumor cells (28). Thus, intracellular Asn levels control uptake of serine, arginine and histidine and thereby controls mTORC1 activation and protein synthesis. Our experiments indicated that the availability of Asn regulated TCR-induced mTORC1 activation, suggesting that Asn may have a similar role as an amino acid exchange factor in T cells. A further, non-exclusive possibility is that Asn deprivation may indirectly influence mTORC1 activation as a consequence of reduced expression of nutrient transporters and uptake of amino acids such as leucine. Consistent with this hypothesis, our data indicated that Asn deprivation resulted in reduced TCR-induced expression of CD98 and capacity for uptake of kynurenine.

Our data have determined that effector CD8^+^ T cell populations have the capacity to function independently of extracellular Asn, as a consequence of ASNS expression. These results also emphasize an additional role for Gln in T cell activation; in the absence of Asn, Gln is required to serve as a nitrogen donor for the synthesis of Asn via ASNS activity (13). Our analyses of *Asns* gene-trap mice strongly suggest that T cell development does not require ASNS expression. It is possible that residual expression of low levels of ASNS in *Asns*^Tm1a^ mice, quantified as <10% of wild-type levels, is sufficient to enable T cell development, or that extracellular Asn levels in the thymus are sufficient for ASNS expression to be redundant. Indeed, the bioavailability of Asn varies between distinct anatomical locations *in vivo* (14). Therefore, there is likely to be varying context dependent requirements for ASNS expression on T cell development, homeostasis and inflammatory responses *in vivo*. Future analysis of the impact of lineage-specific deletion of ASNS in the T cell lineage will resolve this outstanding question. Nonetheless, our data show that the low levels of ASNS expression in peripheral *Asns*^Tm1a^ T cells are insufficient to maintain TCR-induced activation in the absence of extracellular Asn.

It is worthy of note that ASNS activity generates both glutamate (Glu) and Asn, and depletes Gln and aspartate (Asp). The extent to which intracellular Glu levels are affected by ASNS-deficiency is not clear. In this regard, Glu is taken up from the extracellular environment by activated T cells, and intracellular levels of both Gln and Glu increase substantially in T cells activated in nutrient replete conditions (8). Nonetheless, our experiments suggested that *Asns*^Tm1a^ T cell responses were not impaired when extracellular Asn was available, suggesting that the impact of ASNS-deficiency on T cell activation was unlikely to be a consequence of reduced Glu levels. It should be noted that these experiments depended on the use of DMEM medium, that is relatively low in glucose (4mM) and was supplemented with dialyzed serum, in order to control Gln and Asn levels. Under such conditions, control T cell responses were suboptimal, therefore it remains possible that ASNS expression is rate-limiting for optimal T cell responses in nutrient-replete conditions.

In summary, our work has shown that activated T cells coordinate nutrient uptake with upregulated expression of key regulators of intracellular amino acid biosynthesis. Elevated ASNS expression via the mTORC1 pathway is part of a transcriptional programme that endows T cells with the capacity to maximise their nutrient resources during effector immune responses.

## Methods

### Mice

*Asns*^*Tm1a(EUCOMM)/Wtsi*^ (referred to as *Asns*^Tm1a^) gene-trap mice were generated by the International Mouse Phenotyping Consortium (IMPC) and were imported to the University of Leeds St. James’s Biomedical Services (SBS) animal facility from MRC Harwell (Harwell, UK). *Asns*^*T*m1a^ mice were backcrossed with in-house C57BL/6J mice; in all experiments, littermates and/or age-matched in-house bred C57BL/6J mice were used as controls. OT-1 *Rag1*^*-/-*^ (29) mice were maintained in the SBS facility. Where applicable, group sizes are indicated by individual data points.

### Cell culture and stimulation

OT-1 TCR transgenic or polyclonal T cells were obtained from lymph nodes and/or spleens of OT-1 *Rag1*^*-/-*^ or *Asns*^Tm1a^ and control mice, respectively. For experiments investigating the role of extracellular amino acids, cells were cultured in Dulbecco’s Modified Eagle Medium (DMEM) containing 4mM glucose (Gibco, Paisley, UK) supplemented with 5% dialyzed FBS (Gibco), penicillin-streptomycin (Gibco), 50 μM 2-mercaptoethanol (Gibco) ± 2 mM L-glutamine ± 300 μM anhydrous L-asparagine (both Sigma Aldrich, Gillingham, UK). Otherwise, cells were cultured in Iscove’s Modified Dulbecco’s Medium (IMDM, Gibco) containing 25mM glucose, 5% FBS, 50 μM 2-mercaptoethanol, 4 mM L-glutamine and 190 μM freebase L-asparagine. OT-1 T cells were stimulated with 10^−8^M SIINFEKL peptides. For generation of OT-1 CTLs, after 2d of culture, cells were washed and cultured for a further 4 days in media containing 20 ng/mL recombinant human IL-2 (Peprotech, London, UK). For analysis of effector cytokine production, T cells were re-stimulated in media containing 10^−9^M SIINFEKL and Brefeldin A (2.5 μg/mL, Sigma Aldrich). For activation of polyclonal T cells, mixed LN cells were activated using plate-bound anti-CD3ε (1.5 μg/mL) (2C11, BioLegend, London, UK) and soluble anti-CD28 (37.51, BioLegend, 1 μg/mL) as indicated, for 2d, then transferred to media containing 20 ng/mL IL-2. In some experiments, CD8^+^ T cells were purified by negative selection and magnetic sorting (Miltenyi Biotech, Bisley, UK), prior to activation as described above. To assess the role of the mTORC1 pathways in regulating ASNS expression, cells were stimulated in the presence or absence of 100 nM rapamycin (Tocris Bioscience).

### Flow Cytometry and Antibodies

The following antibodies were used: CD4-allophycocyanin (APC) or - brilliant violet 421 (BV421) (Clone GK1.5), CD8β-phycoerythrin-Cy7 (PE-Cy7) (Clone YTS156.7.7), CD25-PE (Clone PC61.5), CD36-PE (Clone HM36), CD44-APC-Cy7 (clone 1M7), CD69-peridinin-chlorophyll-protein Cy5.5 (PerCP Cy5.5) (Clone H1.2F3), CD71-fluoroscein isothyocyanate (FITC) (Clone RI7217), CD98-AlexaFluor 647 (AF647) (Clone 4F2), TCRαβ-FITC (clone H57-597), IFNγ-AF488 (clone XMG1.2), TNF-PerCP Cy5.5 (clone MP6-XT22), granzyme B-pacific blue (PB) (clone GB11), Tbet-PE (clone 4B10) (all BioLegend), eomesodermin-AF488 (Dan11mag) (eBioscience). Unconjugated anti-rpS6 phospho-Ser240/244 (Clone D68F8, Cell Signaling Technology) was counterstained with goat anti-rabbit secondary reagents (Molecular Probes). Live/dead aqua and Zombie NIR dyes were from Life Technologies and BioLegend, respectively. For intracellular staining, cells were fixed in Phosflow fix buffer (BD Pharmingen) or eBioscience FoxP3 fix/permeabilization buffers prior to staining in permeabilization buffers. For analysis of protein synthesis, cells were labelled with O-propargyl-puromycin (OPP, Stratech Scientific, 20μM) and the Click-iT Plus AF488 picolyl azide kit (Life Technologies) according to manufacturers’ instructions. Cycloheximide (Sigma Aldrich, 100μg/ml) was added 15 min prior to labelling as a negative control. For analysis of nutrient uptake, T cells were cultured with 2-deoxy-2-((7-nitro-2, 1, 3-benzooxadiazol-4-yl)amino)-glucose (2-NBDG) (Abcam 50μM, 1h), 4, 4-difluoro-5, 7-dimethyl-4-bora-3a, 4a-diaza-s-indacene-3-hexadecanoic acid (BODIPY™-C16, Invitrogen, 1μM, 30min) or kynurenine (Sigma Aldrich, 800μM in HBSS buffer for 4min), and were washed 3 times in PBS. For BODIPY and Kynurenine uptake, cells were counterstained with Zombie NIR prior to FACS analysis. Samples were acquired with LSR II (Becton Dickinson) or Cytoflex LX (Beckman Coulter) flow cytometers and data were analysed using FlowJo software (Treestar).

### ELISA

To prevent consumption of secreted IL-2, CD25 blocking antibodies (Clone 3C7, BioLegend) were added to culture media prior to T cell stimulation. Levels of cytokines in culture supernatants were assessed using mouse IL-2 and IFN-gamma DuoSet ELISA kits (R&D Systems), according to manufacturers’ instructions.

### Metabolic analyses

ECAR and OCR values were acquired using a Seahorse XFe96 Analyzer, as described previously (30). Briefly, stimulated T cells were plated (10^5^ viable cells/well) in Cell-Tak (Corning) coated Seahorse Analyzer XFe96 culture plates in Seahorse assay media supplemented with glucose (10 mM), glutamine (2 mM) and pyruvate (1 mM). Data were collected in Wave software and analysed using Graphpad Prism.

### ATP assay

T cells were stimulated as indicated in Figure legends. For the measurement of cellular ATP levels, a luminescence-based kit (Abcam, ab113949) was used according to manufacturer’s instructions.

### Western blotting

T cells were stimulated as indicated in Figure legends and cell lysates prepared in RIPA buffer. Protein concentration of lysates was assessed by Bradford Assay (Thermofisher) and 20 μg of protein per sample loaded on polyacrylamide gels. Samples were separated by SDS-PAGE and proteins transferred to nitrocellulose membranes (BioRad). Membranes were blocked in LI-COR blocking buffer (LI-COR Biosciences). Primary antibodies (rabbit anti-ASNS – HPA029318, Atlas Antibodies, mouse anti-β actin, Clone AC-15, Sigma Aldrich) were diluted in LI-COR blocking buffer and detected using goat anti-rabbit-AF680 and goat anti-mouse-AF790 secondary reagents (Molecular Probes). Protein bands were detected and quantified using a Li-COR Odyssey Imaging System.

### Statistical Analysis

Two-tailed Student’s *t-*tests, Mann-Whitney test, and one- or two-way ANOVA with Tukey’s or Sidak’s multiple comparison tests were performed using Graphpad Prism software. P values or adjusted p values < 0.05 were considered to be significant. In graphs, bars represent mean values, dots represent values from replicate samples and error bars represent SDs. Numbers of experimental and/or technical replicates are described in Figure legends.

### Study Approval

OT-1 and *Asns*^*Tm1a*^ mice were bred in accordance with a United Kingdom Home Office Project Licence (PPL-PDAD2D507) and the regulation of the University of Leeds Animal Welfare and Ethics Review Board.

## Author contributions

HCH designed research, performed experiments, acquired and analysed data. RJB and LS performed experiments, acquired and analysed data. RJS designed research, performed experiments, acquired and analysed data and wrote the manuscript. All authors read and discussed manuscript drafts.

## Acknowledgements

We are grateful to G. Doody and G. Cook (University of Leeds) for helpful discussions and comments on manuscript drafts. This work was supported by grants from Cancer Research UK (23269) and UK Medical Research Council (MR/P026206/1) awarded to RJS. HCH is supported by a University of Leeds PhD scholarship.

## References

1. Sinclair LV, Rolf J, Emslie E, Shi YB, Taylor PM, and Cantrell DA. Control of amino-acid transport by antigen receptors coordinates the metabolic reprogramming essential for T cell differentiation. Nat Immunol. 2013;14(5):500–8.

2. Nakaya M, Xiao Y, Zhou X, Chang JH, Chang M, Cheng X, et al. Inflammatory T cell responses rely on amino acid transporter ASCT2 facilitation of glutamine uptake and mTORC1 kinase activation. Immunity. 2014;40(5):692–705.

3. Johnson MO, Wolf MM, Madden MZ, Andrejeva G, Sugiura A, Contreras DC, et al. Distinct Regulation of Th17 and Th1 Cell Differentiation by Glutaminase-Dependent Metabolism. Cell. 2018;175(7):1780–95 e19.

4. Ananieva EA, Powell JD, and Hutson SM. Leucine Metabolism in T Cell Activation: mTOR Signaling and Beyond. Adv Nutr. 2016;7(4):798S–805S.

5. Wei J, Raynor J, Nguyen TL, and Chi H. Nutrient and Metabolic Sensing in T Cell Responses. Front Immunol. 2017;8:247.

6. Mellor AL, and Munn DH. Tryptophan catabolism and T-cell tolerance: immunosuppression by starvation? Immunol Today. 1999;20(10):469–73.

7. Sinclair LV, Howden AJ, Brenes A, Spinelli L, Hukelmann JL, Macintyre AN, et al. Antigen receptor control of methionine metabolism in T cells. Elife. 2019;8.

8. Geiger R, Rieckmann JC, Wolf T, Basso C, Feng Y, Fuhrer T, et al. L-Arginine Modulates T Cell Metabolism and Enhances Survival and Anti-tumor Activity. Cell. 2016;167(3):829–42 e13.

9. Ma EH, Bantug G, Griss T, Condotta S, Johnson RM, Samborska B, et al. Serine Is an Essential Metabolite for Effector T Cell Expansion. Cell Metab. 2017;25(2):345–57.

10. Klysz D, Tai X, Robert PA, Craveiro M, Cretenet G, Oburoglu L, et al. Glutamine-dependent alpha-ketoglutarate production regulates the balance between T helper 1 cell and regulatory T cell generation. Sci Signal. 2015;8(396):ra97.

11. Mak TW, Grusdat M, Duncan GS, Dostert C, Nonnenmacher Y, Cox M, et al. Glutathione Primes T Cell Metabolism for Inflammation. Immunity. 2017;46(4):675–89.

12. Swamy M, Pathak S, Grzes KM, Damerow S, Sinclair LV, van Aalten DM, et al. Glucose and glutamine fuel protein O-GlcNAcylation to control T cell self-renewal and malignancy. Nat Immunol. 2016;17(6):712–20.

13. Lomelino CL, Andring JT, McKenna R, and Kilberg MS. Asparagine synthetase: Function, structure, and role in disease. J Biol Chem. 2017;292(49):19952–8.

14. Knott SRV, Wagenblast E, Khan S, Kim SY, Soto M, Wagner M, et al. Asparagine bioavailability governs metastasis in a model of breast cancer. Nature. 2018;554(7692):378–81.

15. Linares JF, Cordes T, Duran A, Reina-Campos M, Valencia T, Ahn CS, et al. ATF4-Induced Metabolic Reprograming Is a Synthetic Vulnerability of the p62-Deficient Tumor Stroma. Cell Metab. 2017;26(6):817–29 e6.

16. Peng T, Golub TR, and Sabatini DM. The immunosuppressant rapamycin mimics a starvation-like signal distinct from amino acid and glucose deprivation. Mol Cell Biol. 2002;22(15):5575–84.

17. Yang X, Xia R, Yue C, Zhai W, Du W, Yang Q, et al. ATF4 Regulates CD4(+) T Cell Immune Responses through Metabolic Reprogramming. Cell Rep. 2018;23(6):1754–66.

18. Kullas AL, McClelland M, Yang HJ, Tam JW, Torres A, Porwollik S, et al. L-asparaginase II produced by Salmonella typhimurium inhibits T cell responses and mediates virulence. Cell Host Microbe. 2012;12(6):791–8.

19. Torres A, Luke JD, Kullas AL, Kapilashrami K, Botbol Y, Koller A, et al. Asparagine deprivation mediated by Salmonella asparaginase causes suppression of activation-induced T cell metabolic reprogramming. J Leukoc Biol. 2016;99(2):387–98.

20. Kafkewitz D, and Bendich A. Enzyme-induced asparagine and glutamine depletion and immune system function. Am J Clin Nutr. 1983;37(6):1025–30.

21. Sinclair LV, Neyens D, Ramsay G, Taylor PM, and Cantrell DA. Single cell analysis of kynurenine and System L amino acid transport in T cells. Nat Commun. 2018;9(1):1981.

22. Salmond RJ, Emery J, Okkenhaug K, and Zamoyska R. MAPK, phosphatidylinositol 3-kinase, and mammalian target of rapamycin pathways converge at the level of ribosomal protein S6 phosphorylation to control metabolic signaling in CD8 T cells. J Immunol. 2009;183(11):7388–97.

23. Salmond RJ. mTOR Regulation of Glycolytic Metabolism in T Cells. Front Cell Dev Biol. 2018;6:122.

24. Ruzzo EK, Capo-Chichi JM, Ben-Zeev B, Chitayat D, Mao H, Pappas AL, et al. Deficiency of asparagine synthetase causes congenital microcephaly and a progressive form of encephalopathy. Neuron. 2013;80(2):429–41.

25. Pavlova NN, Hui S, Ghergurovich JM, Fan J, Intlekofer AM, White RM, et al. As Extracellular Glutamine Levels Decline, Asparagine Becomes an Essential Amino Acid. Cell Metab. 2018;27(2):428–38 e5.

26. Zhang J, Fan J, Venneti S, Cross JR, Takagi T, Bhinder B, et al. Asparagine plays a critical role in regulating cellular adaptation to glutamine depletion. Mol Cell. 2014;56(2):205–18.

27. Loayza-Puch F, Rooijers K, Buil LC, Zijlstra J, Oude Vrielink JF, Lopes R, et al. Tumour-specific proline vulnerability uncovered by differential ribosome codon reading. Nature. 2016;530(7591):490–4.

28. Krall AS, Xu S, Graeber TG, Braas D, and Christofk HR. Asparagine promotes cancer cell proliferation through use as an amino acid exchange factor. Nat Commun. 2016;7:11457.

29. Hogquist KA, Jameson SC, Heath WR, Howard JL, Bevan MJ, and Carbone FR. T cell receptor antagonist peptides induce positive selection. Cell. 1994;76(1):17–27.

30. Brownlie RJ, Wright D, Zamoyska R, and Salmond RJ. Deletion of PTPN22 improves effector and memory CD8+ T cell responses to tumors. JCI Insight. 2019;5.

