## Supplemental Figs 1 and 2 for "Coordination of asparagine uptake and asparagine synthetase expression is required for T cell activation"

Supplemental Figures 1, 2

A

B

C

*+ IL-2*

*No IL-2*

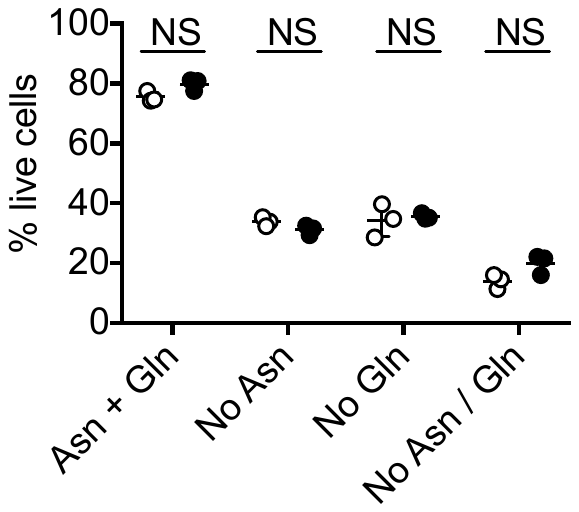

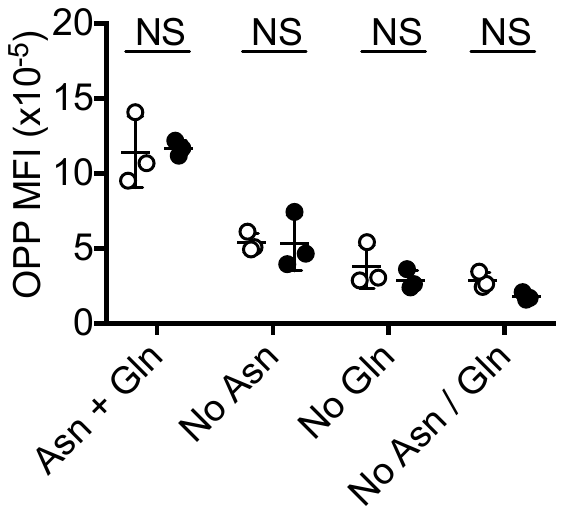

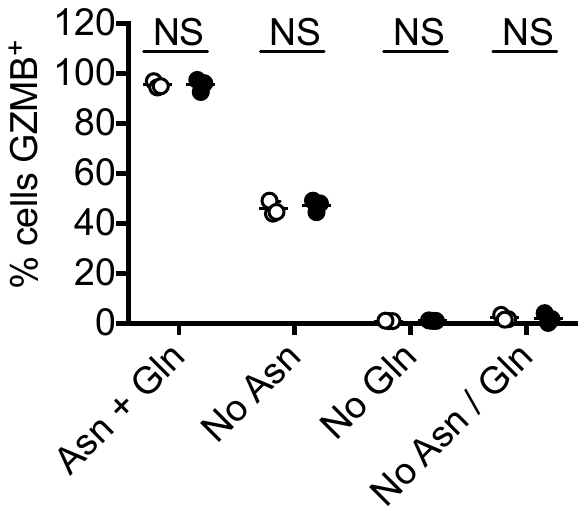

**Supplemental Figure 1: Exogenous IL-2 does not rescue T cell activation in amino acid-deprived conditions**

OT-1 T cells were activated for 24h (A, B) or 48h (C) in DMEM ± Asn and/or Gln, in the presence or absence of 1ng/mL recombinant IL-2. Proportions of live cells (A) were determined by live-dead aqua dye exclusion and FACS. Levels of protein synthesis were assessed by incorporation of OPP, Click chemistry labelling and FACS (B). Intracellular granzyme B expression was assessed by FACS (C). NS – not significant as determined by 2-way ANOVA.

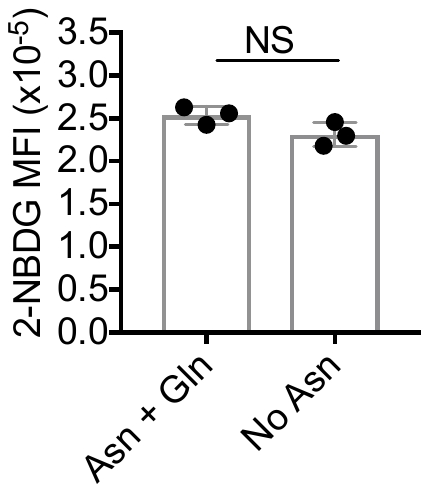

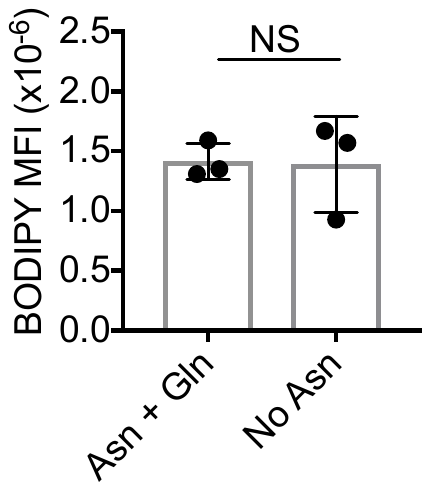

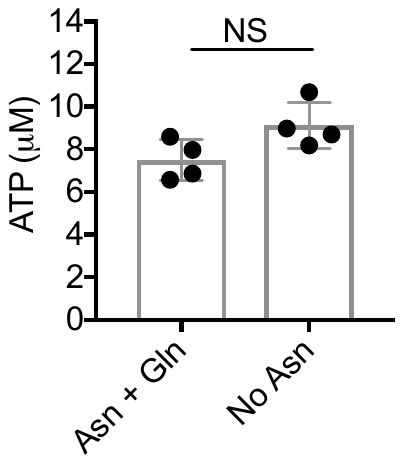

A

B

C

**Supplemental Figure 2: T cells lose dependence upon extracellular Asn upon prolonged activation**

OT-1 T cells were activated for 72h with SIINFEKL peptide in DMEM ± Asn. Uptake of fluorescent 2-NB-d-glucose (A) and BODIPY^TM^-C16 (B) was assessed by FACS. Data shown are mean fluorescence intensities (MFI). (C) Cellular ATP levels were assessed using a luminescent assay (C). NS – not significant, as assessed by Mann-Whitney test.
